# Microbial Risk Score for Capturing Microbial Characteristics, Integrating Multi-omics Data, and Predicting Disease Risk

**DOI:** 10.1101/2022.06.07.495127

**Authors:** Chan Wang, Leopoldo N. Segal, Jiyuan Hu, Boyan Zhou, Richard Hayes, Jiyoung Ahn, Huilin Li

## Abstract

**Background:** With the rapid accumulation of microbiome-wide association studies, a great amount of microbiome data are available to study the microbiome’s role in human disease and advance the microbiome’s potential use for disease prediction. However, the unique features of microbiome data hinder its utility for disease prediction.

**Methods:** Motivated from the polygenic risk score framework, we propose a microbial risk score (MRS) framework to aggregate the complicated microbial profile into a summarized risk score that can be used to measure and predict disease susceptibility. Specifically, the MRS algorithm involves two steps: 1) identifying a sub-community consisting of the signature microbial taxa associated with disease, and 2) integrating the identified microbial taxa into a continuous score. The first step is carried out using the existing sophisticated microbial association tests and pruning and thresholding method in the discovery samples. The second step constructs a community-based MRS by calculating alpha diversity on the identified sub-community in the validation samples. Moreover, we propose a multi-omics data integration method by jointly modeling the proposed MRS and other risk scores constructed from other omics data in disease prediction.

**Results:** Through three comprehensive real data analyses using the NYU Langone Health COVID-19 cohort, the gut microbiome health index (GMHI) multi-study cohort, and a large type 1 diabetes cohort separately, we exhibit and evaluate the utility of the proposed MRS framework for disease prediction and multi-omics data integration. In addition, the disease-specific MRSs for colorectal adenoma, colorectal cancer, Crohn’s disease, and rheumatoid arthritis based on the relative abundances of 5, 6, 12, and 6 microbial taxa respectively are created and validated using the GMHI multi-study cohort. Especially, Crohn’s disease MRS achieves AUCs of 0.88 ([0.85-0.91]) and 0.86 ([0.78-0.95]) in the discovery and validation cohorts, respectively.

**Conclusions:** The proposed MRS framework sheds light on the utility of the microbiome data for disease prediction and multi-omics integration, and provides great potential in understanding the microbiome’s role in disease diagnosis and prognosis.

## Background

Recent microbiome-wide association studies (MWASs) have uncovered that microbiome plays a crucial role in human health and disease [1–4], with linkage of microbiota dysbiosis to a variety of complex diseases, including diabetes, cardiovascular and mental disease, and cancer [5–12]. These studies provide great opportunities to study microbiome’s role in disease prediction, which, however, is challenging because of its unique data structure.

Rapid advances in high-throughput sequencing technologies identify diverse microorganisms in a single sample by targeted sequencing of their unique 16S rRNA gene, or shotgun sequencing of the collective genomes of all microbes. For 16S rRNA sequencing data, QIIME 2 [13] is commonly used to assign the sequencing reads to amplicon sequence variants or clustered operational taxonomic units based on the similarity of sequences. For shotgun sequencing data, MetaPhlAn [14] or StrainPhlAn [15] can be used to map the sequencing reads to species/strains against a reduced set of clade-specific marker sequences. Either method produces the count or relative abundance table which typically contain hundreds to thousands of taxonomic or functional features, i.e. microbiome data are high-dimensional, especially compared to the available number of samples in most existing studies. In addition, these feature tables are usually sparse with excessive zero counts, compositional with a sum constrained to a constant, and heterogeneous with a phylogenetic tree structure to reveal the evolutionary relationship among the taxa. How to deal with these unique characteristics of microbiome data and effectively utilize them in predicting disease risk is challenging and needs comprehensive explorations and validations.

Polygenic risk score (PRS), a continuous score of an individual’s genetic liability to a complex disease or phenotype, has become more routine and powerful in current genomic research [16, 17]. PRS aggregates the results from genome-wide association studies (GWASs) and is defined as the sum of risk alleles linked to a phenotype of interest weighted by the corresponding effect sizes. The construction of PRS involves two key steps: determining the risk alleles and their effect sizes based on discovery samples or published GWASs, and calculating the PRS for each subject in the target population. The PRS framework motivates us to construct a similar microbial risk score (MRS) to summarize the disease-specific microbial profile in the increasing large-scale population-based microbiome studies [11, 18, 19] and to investigate its potential in disease prediction. However, microbiome’s unique community features make MWASs differ from GWASs. First, the microbiota is a complex ecosystem, whose dynamics are driven by the interactions among microbes and between microbes and their host. The link between this complex ecosystem and disease process often involves interwoven mechanisms [20]. Further, the microbiota is composed of various sub-communities related to different traits [21, 22], and its influence on disease development may act at the community rather than the single-microbe level [23]. Thus, it is less informative or efficient to simply define MRS as the weighted sum of the relative abundances of the associated microbes. Instead, we propose a community-based MRS by calculating alpha diversity on a sub-community with member taxa identified as being associated with the study trait. Alpha diversity is the diversity in a single ecosystem or sample with respect to its richness, evenness, or both characteristics [24, 25]. Several indices, including Observed, Simpson, Shannon, and Faith’s phylogenetic diversity (PD), have been extensively used to characterize microbial community. With the NYU Langone Health (NYULH) COVID-19 cohort [26] and the gut microbiome health index (GMHI) multi-study cohort [27], we propose and validate a few MRSs on COVID-19, colorectal adenoma (CA), colorectal cancer (CC), Crohn’s disease (CD), and rheumatoid arthritis (RA) to exhibit the utility of the proposed MRS framework.

With the recent advances in the next-generation sequencing and mass spectrometry, there is a growing need for the ability to merge biological features to study an ecosystem as a whole. Aspects such as the metagenome, metatranscriptome, host genome, host gene expression, and metabolome each provides a snapshot of one level of regulation in a system. The proposed MRS framework provides a simple and interpretable approach to integrate the microbial profiles with other biological omics data and elucidate the microbial interactions with other omics datasets in the disease prediction. We use the NYULH COVID-19 cohort, which characterized the lung microbiome in a large prospective cohort of critically ill patients with SARS-CoV-2 infection who required invasive mechanical ventilation, to illustrate, evaluate, and validate the proposed MRS and its integrations with other omics data in the prediction for COVID-19 mortality. In addition, we elucidate the join effect of MRS and PRS on T1D risk stratification using the Environmental Determinants of Diabetes in the Young (TEDDY) study (https://teddy.epi.usf.edu/) [28–30].

## Methods

### MRS framework

#### MRS workflow

We propose a microbial risk score framework to convert the high-dimensional microbiome data into a summarized risk score that can measure and predict disease susceptibility. As illustrated in Figure 1, with the ready-for-downstream-analysis microbial data, the microbial risk score algorithm involves two key steps: 1) to identify a sub-community consisting of the signature microbial taxa associated with disease, and 2) to integrate the identified microbial taxa into a continuous score.

**Figure 1.**
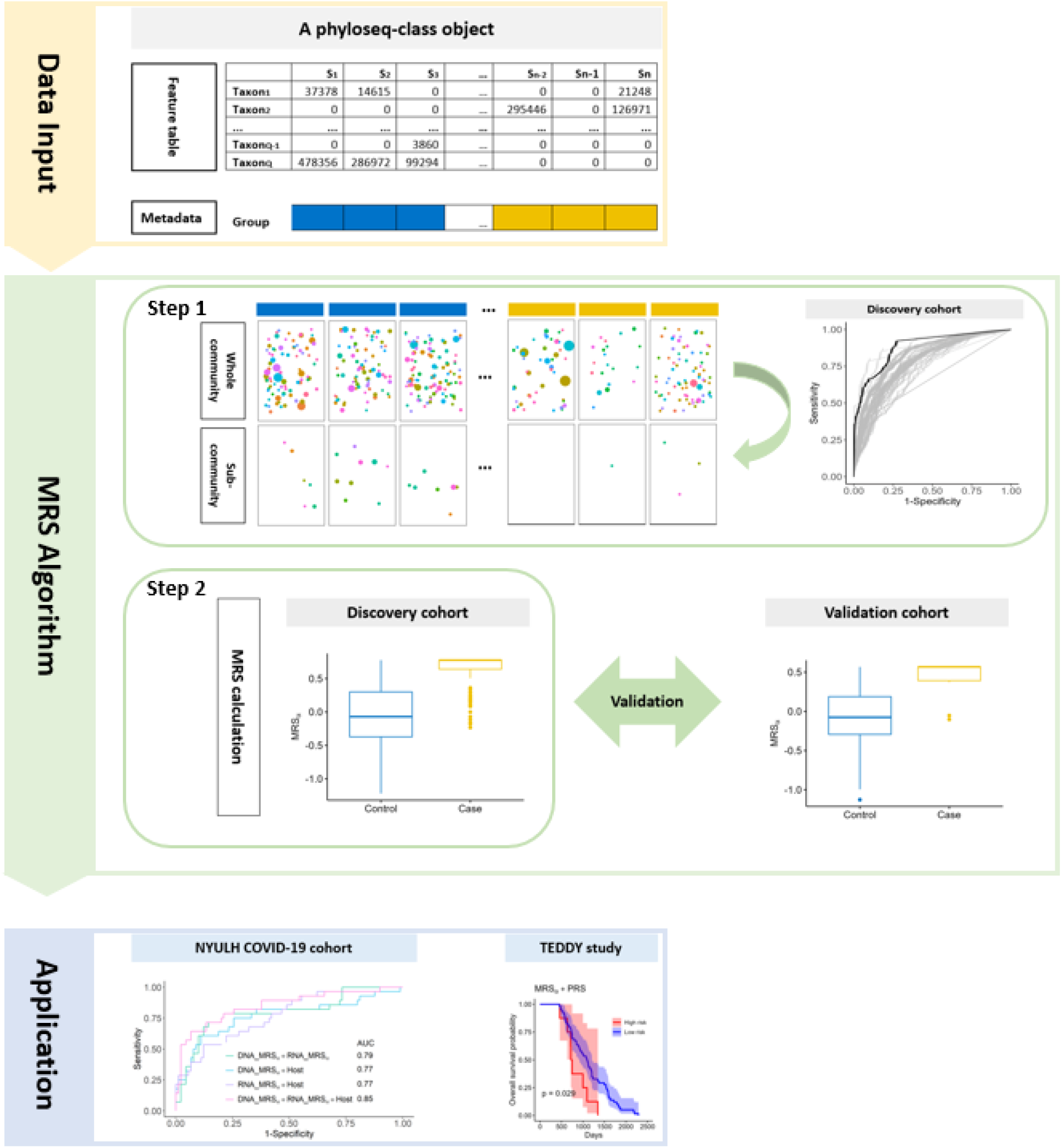
The workflow of the microbial risk score (MRS) framework. Data Input: a phyloseq-class object is needed, which consists of a feature table (observed count table), a sample metadata, a taxonomy table (optional), and a phylogenetic tree (optional). MRS Algorithm has two steps: Step 1 is to identify a sub-community consisting of the signature microbial taxa with the P+T method and AUC evaluation in the discovery cohort. The black ROC curve which has the largest AUC determines the optimal *p-*value cutoff. Step 2 is to integrate the identified microbial taxa into a continuous score, i.e., calculate the MRS value for each sample by calculating the diversity of the identified sub-community with the Shannon index. In addition, the constructed MRS is independently validated in the validation cohort. Application: In this manuscript, we perform multi-omics data integration for disease prediction by jointly modeling the proposed MRS and other risk scores constructed from other omics data in two real data cohorts.

#### Microbial signature identification

We propose to employ the existing sophisticated microbial association tests [1, 3, 4, 31-33] to identify microbial taxa associated with disease using the discovery samples. Great amount of abundance-based methods examining the difference of microbial abundance directly, which is also called differential abundance (DA) analysis [31–39] have been proposed recently. Based on the results in two recent benchmarking works [32, 33], ANCOM-BC (Analysis of compositions of microbiomes with bias correction) [31] is one of the top-performing methods and has been widely used in microbiome research.

ANCOM-BC [31] models the observed abundances using an offset-based log-linear model, in which the offset term is sample-specific to account for sampling fraction. We use it as the default microbial association test to identify the candidate taxa in the first step of our microbial risk score algorithm. Considering developing novel differential abundance test is still an active area of research, in the Discussion section, we discuss the performance of the proposed MRS framework with other DA tests.

In addition to the above-mentioned statistical methods, a variety of machine learning (ML) techniques have been applied in microbiome studies for microbial feature selection, biomarker identification, disease prediction and classification, as recently reviewed [40]. As an example, Gou et al. [41] defined an MRS with the microbiome features selected by the Light Gradient Boosting Machine method [42] and examined its association with type 2 diabetes (T2D) as well as T2D-related traits. Despite the visible contributions of characterizing the microbial profiles and uncovering the relationship between microbiome and disease, the applications of ML methods including traditional methods and deep learning techniques in the microbiome studies share several drawbacks [40]. One is that most ML methods input all available microbial features into the model to determine the final output solely based on algorithms, without considering the inherent structure of microbiome data, such as compositionality and zero inflation. Another unavoidable drawback of ML methods is the model instability in the relatively small-scale biomedical human studies [43]. Because the nature of ML algorithms is to learn the pattern by training the data, they usually require a large sample size to reach stable results, especially for the algorithms involving various parameters or various layers that need to be trained via cross-validation (CV). Given these common pitfalls and relatively small sample size in biomedical studies due to the high cost of patients’ in-person visit, sample collection and sequencing, ML’s application in microbiome research may provide inexplicable results and even lead to the loss of statistical power. With the NYULH COVID-19 cohort example, we illustrate the inefficient utility of ML methods in analyzing the microbiome data compared to the proposed MRS method. The details are reported in the Results section.

#### Sub-community determination

Pruning and thresholding (P+T) method is a heuristic approach commonly used in PRS studies for identification of genetic variants based on an empirically determined *p*-value threshold [44]. We propose to use P+T method to determine the final candidate microbial taxa in discovery cohort. Specifically, we calculated a series of MRSs proposed below using the nested sets of microbial taxa with the increasingly relaxed significance thresholds. We set the final threshold at the value that produced the largest area under the receiver operating characteristic (ROC) curve (AUC). All the taxa whose p-values are less than the final threshold form a disease or trait specific sub-community. If there is only one dataset available, CV will be used to determine the sub-community along with P+T method. More details are provided in the Results section.

#### MRS calculation

We propose an MRS, denoted by MRS_*α*_, which is defined as the alpha diversity of the sub-community consisting of the identified candidate taxa. Alpha diversity is the diversity in a single ecosystem or sample with respect to its richness, evenness, or both characteristics [24, 25]. The core concept of alpha diversity index in biology is to find the effective number of elements of a system to measure its complexity or diversity [45]. Note that multiple alpha diversity indices are available. Some measure species richness such as observed index, Chao1, and ACE. PD is a phylogenetic metric which is defined as the sum of the lengths of all those branches on the tree that span the members of the set. Simpson index is a dominance index as it gives more weight to the common or dominant species and does not account for species richness. While Shannon index is an information statistic index (entropy) which accounts for both species richness and its evenness in a community or sample, and it has a unique ability to weigh taxa by their frequency, without disproportionately favoring either rare or common elements. As the most popular and accepted index for diversity [46], we adopt Shannon index in the proposed MRS_*α*_. Other indices are also investigated in the Discussion section and included in the MRS framework (MRS R package).

Suppose there are *n* samples (each sample represents one ecosystem or microbial community) and *Q* taxa. Let *M_ij_* be the relative abundance of the *j*th taxon in the *i*th sample with the constraint 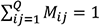, *i* = 1, …, *n*, and *j* = 1, …, *Q*. Assume *p* (<*Q*) taxa are identified as a sub-community to construct MRSs. Without loss of generality, we assume that the first *p* taxa are the identified candidate taxa.

For the *i*th sample, its MRS_*α*_ is calculated as 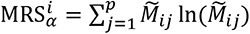 where 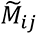 is relative abundance of the *j*th identified candidate taxon within the sub-community for the *i*th sample 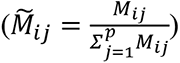.

MRS_*α*_ is constructed based on the Shannon index [24, 25] without the negative sign, so that the smaller is MRS_*α*_, the healthier is the microbial community [47]. As a comparison, we also derive a standard MRS as an analogy to PRS, denoted by MRS_*S*_. It is a (weighted) sum of relative abundances of the identified candidate taxa as 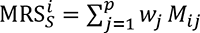, where *w*_*j*_ is the weight for the *j*th taxon. We propose two sets of weights: all weights are equal to 1 (denoted by 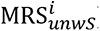); and the weights are the effect sizes estimated from the training or discovery samples by certain microbial association method (denoted by 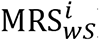). Noticeably, MRS_*α*_ integrates *p* identified taxa as a community by measuring its diversity. While, MRS_*S*_ focuses on the additive effect of the identified taxa and doesn’t account for the microbial community feature.

#### Validation

The proposed MRSs need to be validated either by external validation or internal validation. Since the GMHI multi-study cohort [27] has independent discovery and validation cohorts, the MRSs are created using the discovery cohort and validated using the validation cohort. For the NYULH COVID-19 [26] and TEDDY studies [28–30], due to the lack of independent additional samples, we employ CV to perform independent internal validation.

### Risk score-based multi-omics data integration

Note that the proposed MRS summarizes a complex microbial profile into a quantifiable score, which provides a fast and flexible way to integrate microbiome data with other omics data to better predict disease risk. Both the NYULH COVID-19 and TEDDY studies contain not only microbial profile data, but also other omics data. We propose to jointly model MRS and other risk scores built on other omics data to further improve the performance of disease prediction. In the COVID study, on one hand, the enrichment of SARS-CoV-2 and some oral commensals in the lower-airway microbiota are associated with poor outcome, and on the other hand, host lower-airway immune phenotypes reveal a failure of adaptive and innate immune response to SARS-CoV-2 among deceased subjects. Jointly modeling these omics profiles can improve the predictive accuracy of mortality. For the TEDDY study, since that genotype data in the regions containing autoimmunity and inflammatory response genes are available, one can build a PRS for each subject using the existing PRS algorithms [48–50]. By combining the PRS and the proposed MRS, we can jointly model the association of genetic and environmental risk in T1D prediction.

### Prediction performance evaluation

With the constructed risk scores from various omics data, one can employ a logistic regression model for the prediction of disease status(binary outcome), or a Cox proportional-hazards model [51] for the prediction of disease onset (survival outcome). Predication performance can be evaluated by AUC for binary outcome or by hazard ratio (HR) for survival outcome. The additive model can be used to integrate multiple risk scores in these two regression models. The interaction terms between scores can be explored further for risk stratification [52], as illustrated in the TEDDY study in Result section.

### NYULH COVID-19 cohort

The NYULH COVID-19 cohort [26] includes a subset of 142 patients with COVID-19, at the NYULH Manhattan campus from March 3 to June 18, 2020, who required invasive mechanical ventilation and underwent bronchoscopy for airway clearance and/or tracheostomy. Among all patients, 108 (76%) survived hospitalization and 34 (24%) died. The study has collected and processed lower-airway samples and performed: a) metagenomic sequencing for bacterial, fungal and DNA viral genomes; and b) metatranscriptome assays for viral, bacterial, fungal, and human transcriptomes and the RNA virome. In addition, comprehensive demographic, longitudinal clinical, and treatment data are available.

### GMHI multi-study cohort

An integrated dataset of 4,347 human stool metagenomics samples (cross-sectional) from 34 published studies (discovery cohort) and an independent dataset of 679 samples from 9 additional studies (validation cohort) are publicly available [27]. Both cohorts consist of healthy subjects and patients with various diseases. Using these two cohorts, Gupta et al. [27] introduced and validated the gut microbiome health index (GMHI) to quantify the likelihood of disease presence based on subject’s gut microbiome data. In both cohorts, they pooled samples from different disease conditions together into one nonhealthy group, and the proposed GMHI exclusively identifies the difference of microbiome profile between healthy and non-healthy samples. After the pre-processing and quality control, there are 2,636 healthy and 1,711 nonhealthy samples in the discovery cohort and 118 healthy and 561 nonhealthy samples in the validation cohort respectively. Among nonhealthy samples, discovery and validation cohorts both have samples from patients with CA, CC, CD, and RA. Sample sizes are shown in Table S1. For microbiome data, there are 313 species and 576 species in the discovery and validation cohorts respectively available for analysis. More details are described in Gupta et al. [27].

### TEDDY cohort

TEDDY is a large-scale prospective study designed to identify the genetic and environmental triggers that cause childhood T1D [28–30]. Children with high genetic risk for islet autoimmunity or T1D were enrolled and multiple biomarkers were assessed longitudinally for prediction of T1D development. A total of 12,005 fecal samples from 903 children, collected from 3 to 46 months of age, were characterized by 16S rRNA sequencing. Of this cohort, 114 children were ascertained to T1D by year 5 [29]. The findings in the previous TEDDY publications [53, 54] focus exclusively on the microbiome profiles, and suggest that the gut microbiome data may have the potential to predict the progression of T1D. In addition to microbiome data and metadata, the TEDDY cohort also includes genomic, longitudinal metabolomic, and host transcriptomic data which together provide opportunity to explore the integrated information from multiple aspects on the pathogenesis of T1D through the multi-omics analysis.

## Results

### Evaluation and validation of MRS framework

#### NYULH COVID-19 cohort

With the same quality control, sequencing process, and filtering criteria described in Sulaiman et al. [26], we analyzed data from 118 patients (28 Deceased and 118 Alive) who had all metagenome, metatranscriptome, and host transcriptome samples. We included 374 taxa in metagenome, 1,149 taxa in metatranscriptome, and 14,697 genes in host transcriptome data for our analyses. We used the binary outcome (Deceased vs. Alive) to illustrate the predictive performance of MRS here.

Figure 2 presents the optimal *p*-value thresholds (0.42, 0.38, and 0.02) used to identify the associated taxa in MRSs (MRS_*α*_, MRS_*wS*_, and MRS_*unwS*_, respectively) using the metagenomic data. The optimal thresholds were determined by P+T method as described in the sub-section “Sub-community determination**”** using the leave-one-out CV. With the optimal *p*-value cutoffs, the community-based MRS_*α*_ has the best performance in predicting deceased/alive status (AUC=0.74), compared to two summation-based standard MRSs: MRS_*wS*_ (AUC=0.72) and MRS_*unwS*_ (AUC=0.70). This reflects that analyzing the microbial profile as a community can characterize more microbial information and work better than analyzing microbes individually. Additionally, MRS_*wS*_ performs better than MRS_*unwS*_, as expected, since MRS_*wS*_ incorporates the strength of the association effects of taxa on the outcome, as well as the microbial relative abundances, while MRS_*unwS*_ is just the summation of the microbial relative abundances from the selected taxa.

**Figure 2.**
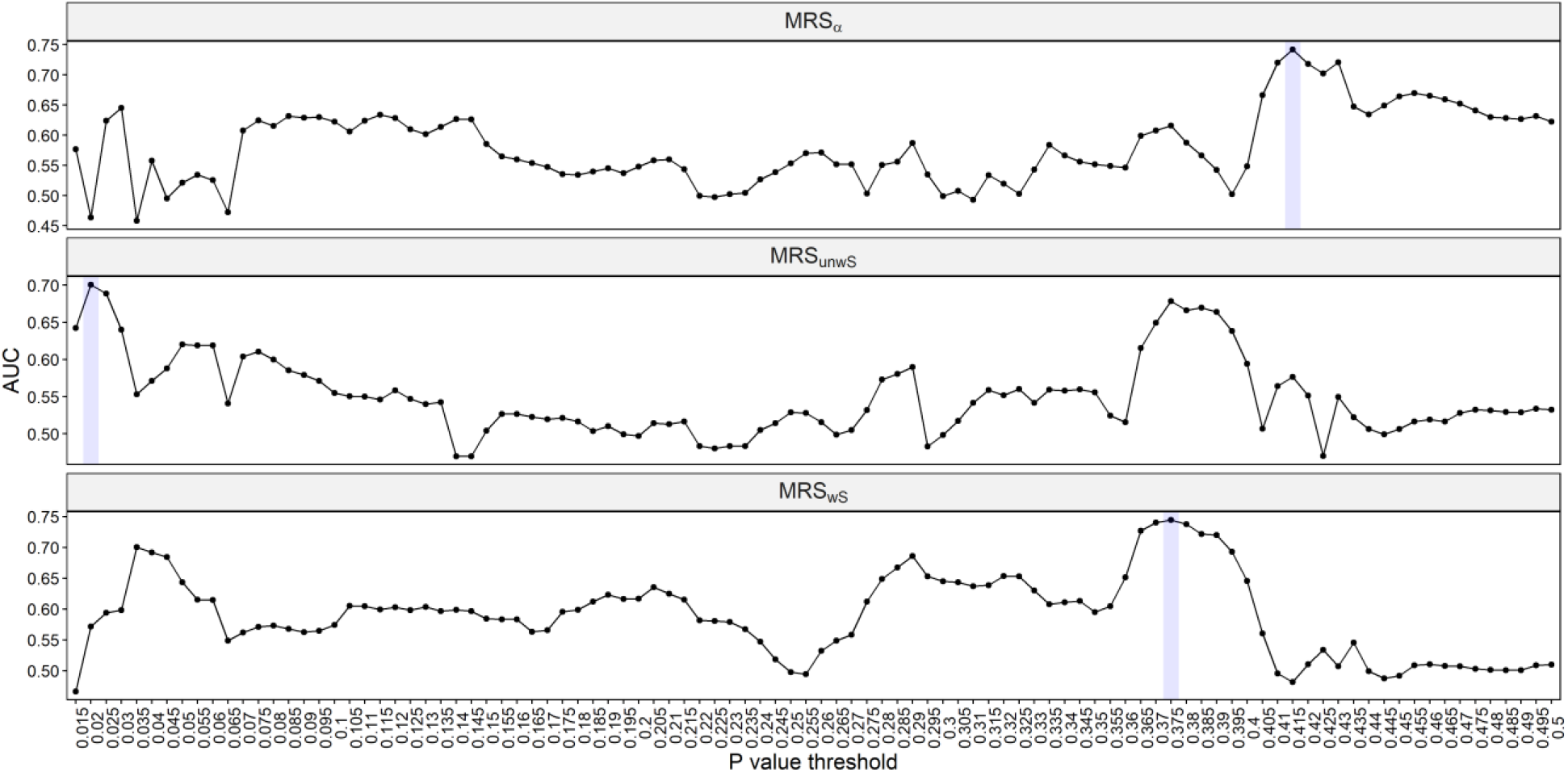
The optimal *p*-value thresholds by P+T method for including taxa in MRS_*α*_, MRS_*wS*_, and MRS_*unwS*_, separately, using the metagenomic data in the NYULH COVID-19 cohort. Specifically, given a cut-off, the taxa with *p*-values less than the cut-off were selected and defined as a sub-community. The *p*-values were obtained by ANCOM-BC method. The leave-one-out CV was used for the predictions. MRS_*α*_: the negative alpha diversity (Shannon index) was calculated for each sample on the selected sub-community; MRS_*wS*_: the weighted sum of relative abundances of the selected taxa with the weights being the coefficients estimated from the ANCOM-BC log-linear model; MRS_*unwS*_: the sum of relative abundances of the selected taxa.

Figure S1 shows prediction performance for various ML algorithms which have been commonly applied in microbiome research [40]. The leave-one-out CV was used for the predictions and the predicted probability for deceased/alive status was used for ROC analysis. All ML algorithms have lower AUCs than the proposed MRS_*α*_. Among these ML algorithms, the ML algorithms based on regularization (Figure S1A) all perform better with higher AUCs, compared to the ML algorithms that have various tuning parameters or layers (Figure S1B). Elastic-net logistic regression and penalized discriminant analysis (regression-based) algorithms have the best prediction performance. On the other hand, ML algorithms were also applied to select the candidate taxa used for the construction of MRS_*α*_ based on the variable importance. The top K features were determined based on leave-one-out CV. Take the elastic-net logistic regression which has the best prediction above for example, the top 30 taxa were ultimately selected to construct MRS_*α*_ with the AUC being the largest based on CV, and its AUC for deceased/alive status prediction is 0.66, which is 11% lower than the AUC of the above MRS_*α*_. The efficiency of ML algorithms is evidently limited due to the small sample size and not being able to take care of the unique features of microbiome data, such as compositionality and zero inflation.

In addition, we checked the prediction performance of the alpha diversity indices on the whole microbial community in terms of AUC. Table S2 reports the AUC values for six common alpha diversity indices in predicting alive and deceased status. All alpha diversity indices have similar prediction performance, with AUC being 0.50 to 0.53, which are much poorer than the proposed MRS_*α*_. Comparisons between MRS_*α*_ and alpha diversity indices underline the significance of identification of the associated taxa in the microbial risk score framework, which condenses the signal by excluding the non-associated taxa and provides full potential for the proposed MRS to measure and predict disease susceptibility.

#### GMHI multi-study cohort

With the discovery and validation cohorts [27], we evaluated and validated the proposed MRS_*α*_ in terms of predictive performance. Specifically, for CA, CC, CD, and RA diseases, respectively, we performed ANCOM-BC to identify candidate species that were differentially abundant between samples from healthy subjects and patients with this disease in the discovery cohort, constructed disease-specific MRS_*α*_ based on the identified species, and performed the independent validation of disease-specific MRS_*α*_ using samples from healthy subjects and patients with the disease in the validation cohort.

Figure 3A presents that AUC values and 95% confidence intervals for MRS_*α*_s to predict healthy and 4 different diseases in discovery and validation cohorts, respectively. Overall, MRS_*α*_s achieve great predictive performance in both discovery (AUCs: 0.60-0.88) and validation (AUCs: 0.68-0.86) cohorts. Notably three MRS_*α*_s (healthy vs. CA, healthy vs. CC, and healthy vs. RA) have higher AUCs in validation cohort, compared to discovery cohort. Among these four disease-specific MRS_*α*_s, MRS_*α*_ specific for CD disease has the best predictive performance (AUC=0.88 in discovery and AUC=0.86 in validation). In addition, different MRS_*α*_s are constructed by different identified taxa. 5, 6, 12, and 6 taxa are used for constructions of MRS_*α*_s for CA, CC, CD, and RA, respectively (Figure 3B; Table S3).

**Figure 3.**
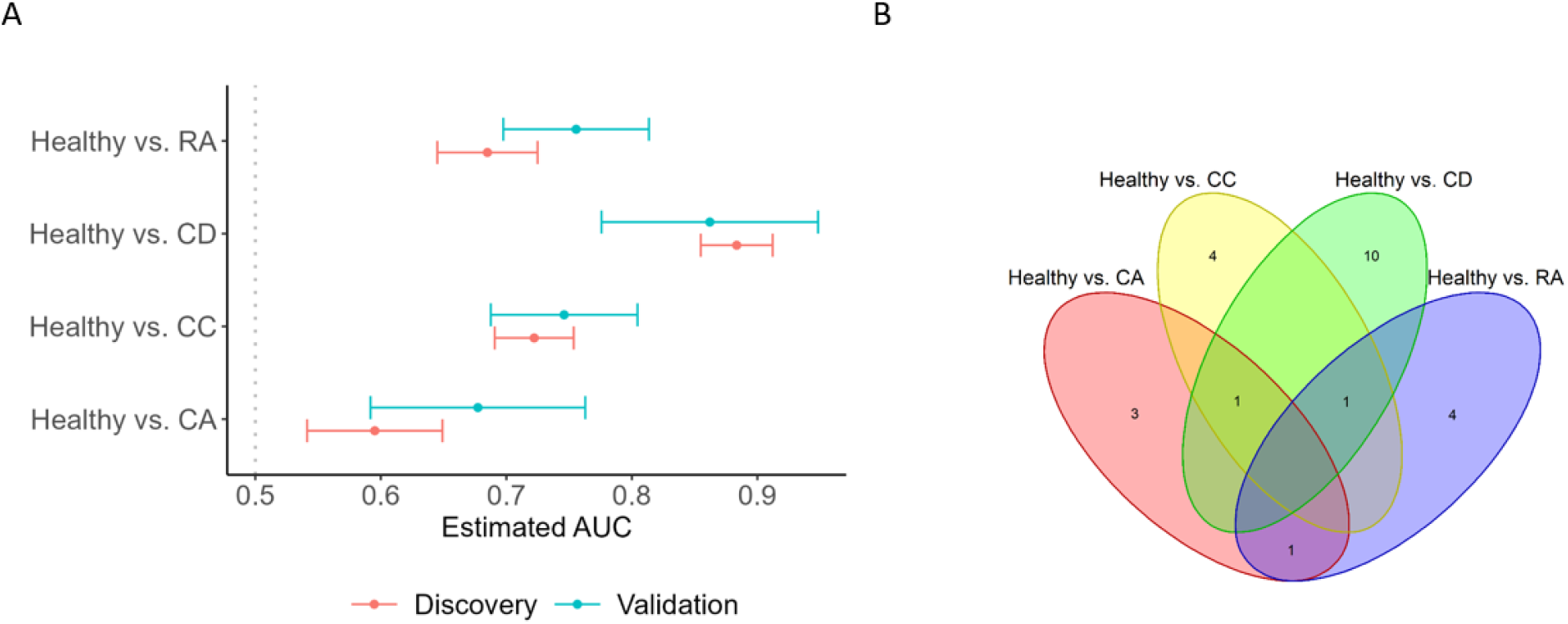
Evaluation of MRS in the discovery and validation cohorts [27]. A: The AUC values and 95% confidence intervals (CIs) for MRS_*α*_s to predict healthy and different disease conditions in discovery and validation cohorts, respectively. B: Venn diagrams of taxa identified in pairwise comparisons of Healthy versus CA, CC, CD, and RA. CA: colorectal adenoma, CC: colorectal cancer, CD: Crohn’s disease, and RA: rheumatoid arthritis.

Several taxa contribute multiple MRS_*α*_s, for example, species *Bifidobacterium angulatum* is involved for constructions of MRS_*α*_s for CA, CC, and RA (Table S3). On the other hand, 21 taxa are disease-specific and exclusively used in one MRS_*α*_ (Table S3). They are differentially abundant in Healthy, CA, CC, CD, and RA samples (Tables S4 and S5). This demonstrates that the proposed MRS framework powerfully improves disease prediction by incorporating the disease-specific microbial profile. This feature makes the proposed MRS framework more crucial in practice, as most research studies aim to identify the microbial taxa specifically playing a role in a certain disease, rather than the generalized disease-associated microbial taxa.

Similar to disease-specific MRS, we also assessed the MRS framework that distinguishes two disease groups, as well as healthy and nonhealthy conditions defined as in the original study [27] in the discovery and validation cohorts, respectively. Figure S2 presents the AUC values and 95% confidence intervals for MRS_*α*_s to classify any two diseases of CA, CC, CD, and RA, and healthy and nonhealthy conditions in discovery and validation cohorts, respectively. Table S3 correspondingly reports which taxa are involved for these MRS_*α*_ calculations, respectively. Again, the MRS framework achieves notable performance. For example, discovery cohort has AUCs of 0.91 and 0.89, meanwhile, validation cohort has AUCs of 0.84 and 0.84, to distinguish CD from RA and CC, respectively. Validation cohort has a relatively lower AUC for classifying CA and RA, due to the small sample size. In terms of healthy vs. nonhealthy prediction, MRS_*α*_ achieves consistently competitive performance but with much fewer species, whose AUCs are 0.7 and 0.71 in discovery and validation cohorts, respectively, compared to GMHI whose AUCs are 0.7 and 0.74 in discovery and validation cohorts, respectively. And the identified 6 species for MRS_*α*_ construction is a subset of 50 microbial species used in GMHI [27].

### Results of risk score-based multi-omics data integration

#### NYULH COVID-19 cohort

In addition to metagenome data, the NYULH COVID-19 cohort has metatranscriptome and host transcriptome data. In the following, we present how to integrate metagenomic, metatranscriptomic, and host transcriptomic datasets using the proposed MRS_*α*_ and the evaluation of different methods. For the metatranscriptomic data, we employed the same MRS algorithm as we described in the Methods section, in terms of determining the *p*-value cutoff, identifying candidate taxa, and constructing the microbial risk score, to construct its MRS_*α*_. In order to differentiate various MRS_*α*_s, we denoted the MRS_*α*_ using the metagenomic and metatranscriptomic data by DNA_MRS_*α*_ and RNA_MRS_*α*_, respectively in the rest of manuscript. For the transcriptomic data, we employed DESeq2 [36] to evaluate the association effects of genes on the deceased/alive status, determined the *p*-value cutoff based on the P+T method, and identified the candidate genes by AUC evaluation. Then we defined the weighted sum of log-transformed counts of the selected candidate genes for each sample as the risk score (denoted as Host), with the weight being 1 if the corresponding logarithmic fold change estimate from DESeq2 was positive, otherwise -1. Computational details are reported in Section S1. Figure 4A shows that the risk scores based on metagenomic, metatranscriptomic, and host transcriptomic data separately have the AUC values of 0.74, 0.69, and 0.63, respectively, in terms of predicting deceased/alive status. Furthermore, the combinations of risk scores from different datasets can obviously improve the predictive performance (Figure 4B) of mortality. The combinations of any two datasets have comparable AUC values and perform similarly. As expected, the integration of all three datasets (DNA_MRS_*α*_ + RNA_MRS_*α*_ + Host) has the highest AUC of 0.85, which yields at least a 15% increase in AUC compared to DNA_MRS_*α*_, RNA_MRS_*α*_, or Host alone.

**Figure 4.**
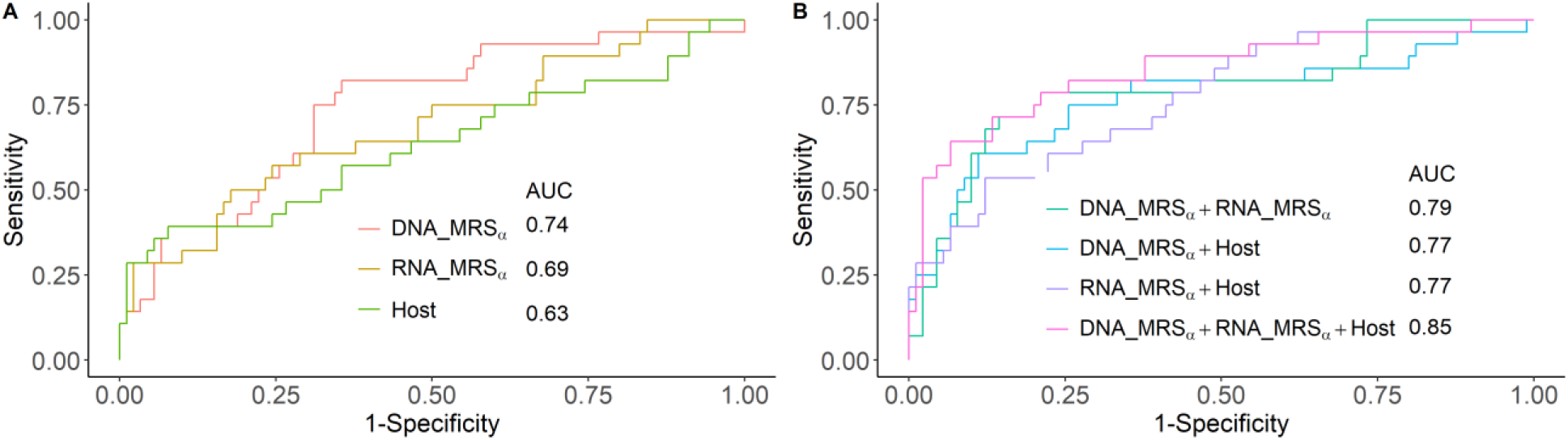
The ROC curves and AUC values for the various risk scores to predict alive and deceased status in the NYULH COVID-19 cohort. A. Predication performance for the individual risk scores constructed based on metagenome (DNA_MRS_*α*_), metatranscriptiome (RNA_MRS_*α*_), and host transcriptome (Host), separately. B. Predication performance based on multiple risk scores using additive model.

In Figure 5, comparing the risk scores between the alive and deceased groups, the deceased group always has a significantly higher average risk score than the alive group, no matter the score was constructed based on a single omics dataset or the integration of different omics datasets (*p*-values<0.05).

**Figure 5.**
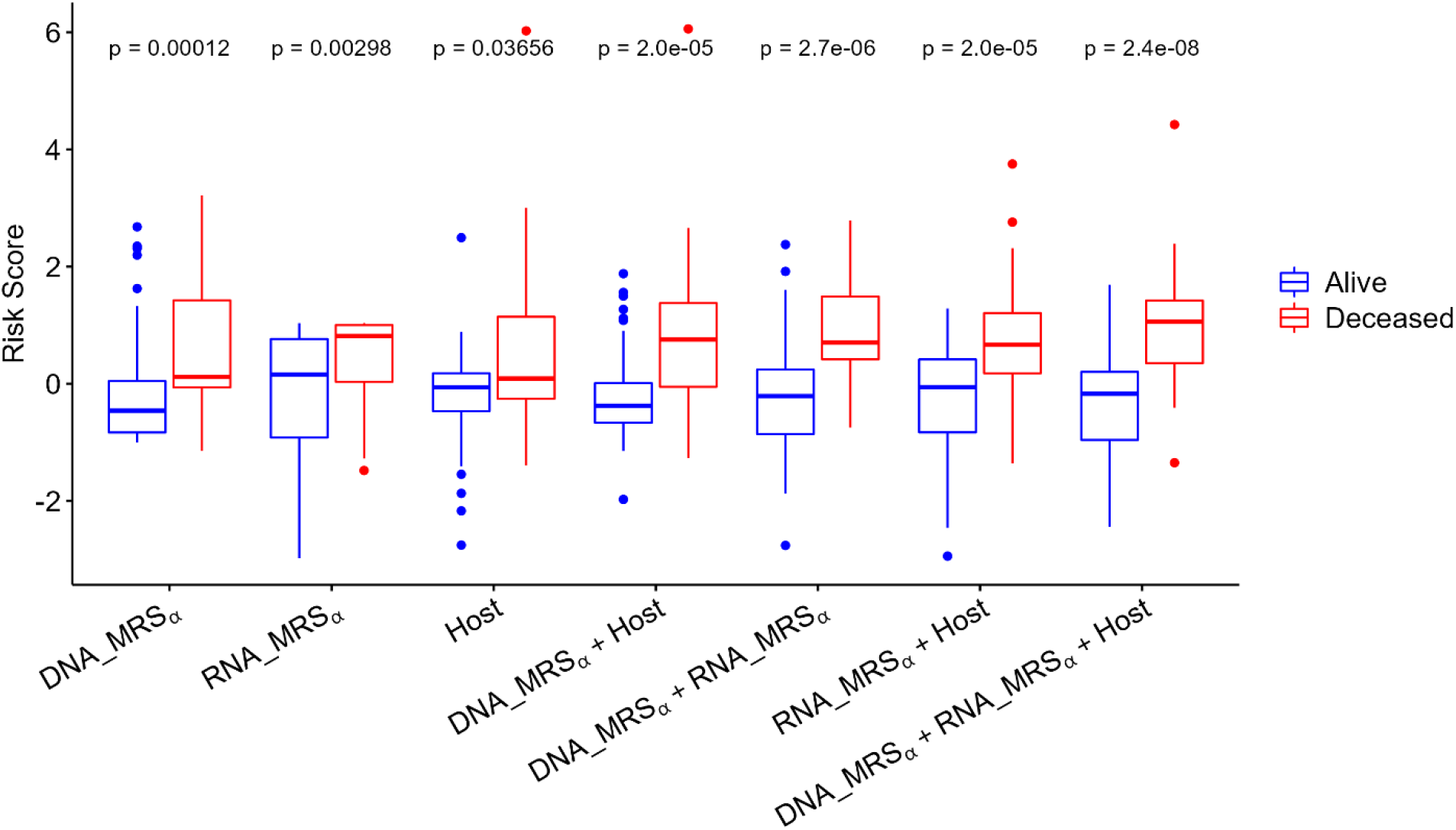
Box plots of the score comparisons between alive and deceased group. All risk scores are standardized among all samples, respectively. The statistical significance on group comparison is evaluated by Wilcoxon signed-rank test.

Figure 6 presents the 2D or 3D scatterplots of risk scores from metagenomics, metatranscriptomic, and host transcriptomic data. The subjects were first classified into “High risk” and “Low risk” groups by each risk score’s mean. We next checked how well these risk classifications can be used to predict disease status by reporting the classification metrics [55]: sensitivity, specificity, accuracy, and F1 score in Table 1. Specifically, the predicted values for the subjects labeled as “High risk” by two risk scores (in Figures 6A-C) or by all three risk scores (in Figure 6D) datasets are “Deceased”, and the predicted values for the subjects labeled as “Low risk” by two risk scores (in Figures 6A-C) or by all three risk scores (in Figure 6D) datasets are “Alive”. From Table 1, we can see that among the combinations of two risk scores for classification, the combination of metagenomic and host transcriptomic risk scores has the highest sensitivity, accuracy and F1 score, but is still inferior to the combination of all three omics risk scores, which identifies the mortality status with 86% sensitivity, 91% specificity, 88% accuracy, and an F1 score of 0.89. In this real study, from different angles, including the AUC in Figure 4, the scatterplots of risk scores in Figure 7, and the test results in Table 1, we show that combining risk scores from metagenomics, metatranscriptomic, and host transcriptomic data increases the predictive accuracy for COVID-19 mortality.

**Table 1.**
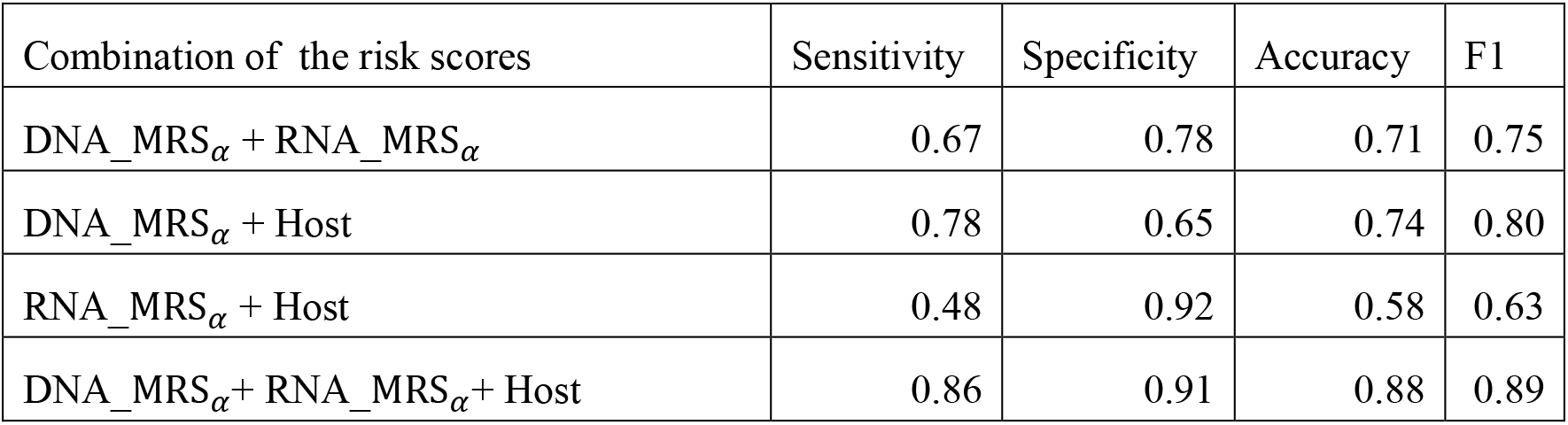
Classification evaluation for subjects having extreme risk categories (labeled as either “High risk” or “Low risk” by both or all three risk scores) in the NYULH COVID-19 cohort.

| Combination of the risk scores | Sensitivity | Specificity | Accuracy | F1 |
| --- | --- | --- | --- | --- |
| DNA_MRS$_{\alpha}$ + RNA_MRS$_{\alpha}$ | 0.67 | 0.78 | 0.71 | 0.75 |
| DNA_MRS$_{\alpha}$ + Host | 0.78 | 0.65 | 0.74 | 0.80 |
| RNA_MRS$_{\alpha}$ + Host | 0.48 | 0.92 | 0.58 | 0.63 |
| DNA_MRS$_{\alpha}$ + RNA_MRS$_{\alpha}$ + Host | 0.86 | 0.91 | 0.88 | 0.89 |

**Figure 6.**
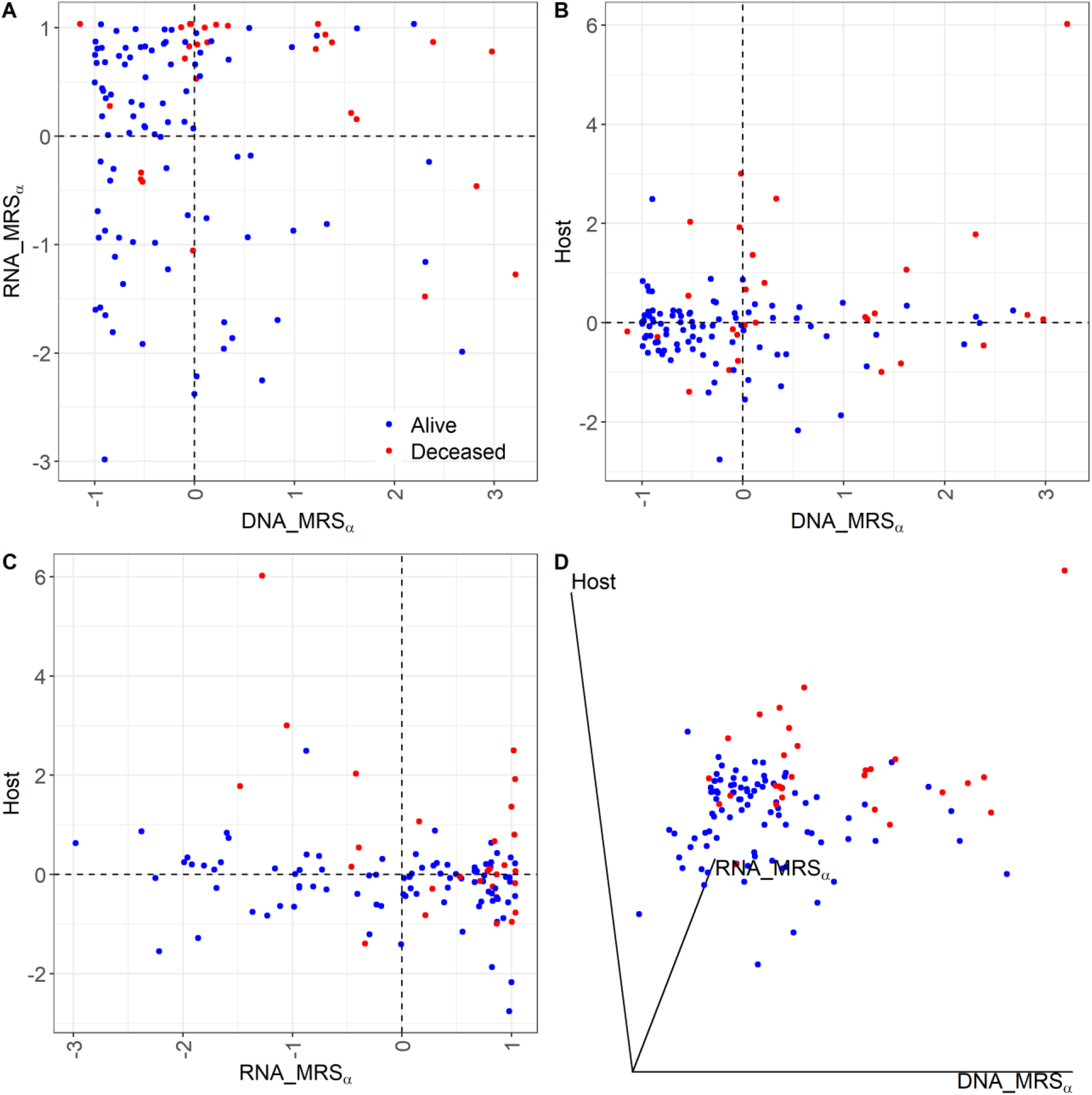
Scatterplots of risk scores based on metagenome, metatranscriptome, and host transcriptome data. A-C: Scatterplots of DNA_ MRS*α* vs RNA_ MRS*α*, DNA_ MRS*α* vs Host, and RNA_ MRS*α* vs Host, respectively. Dotted line denotes the mean of the corresponding risk score across all subjects. D: 3D scatterplot of DNA_MRS_*α*_vs RNA_MRS_*α*_vs Host.

**Figure 7.**
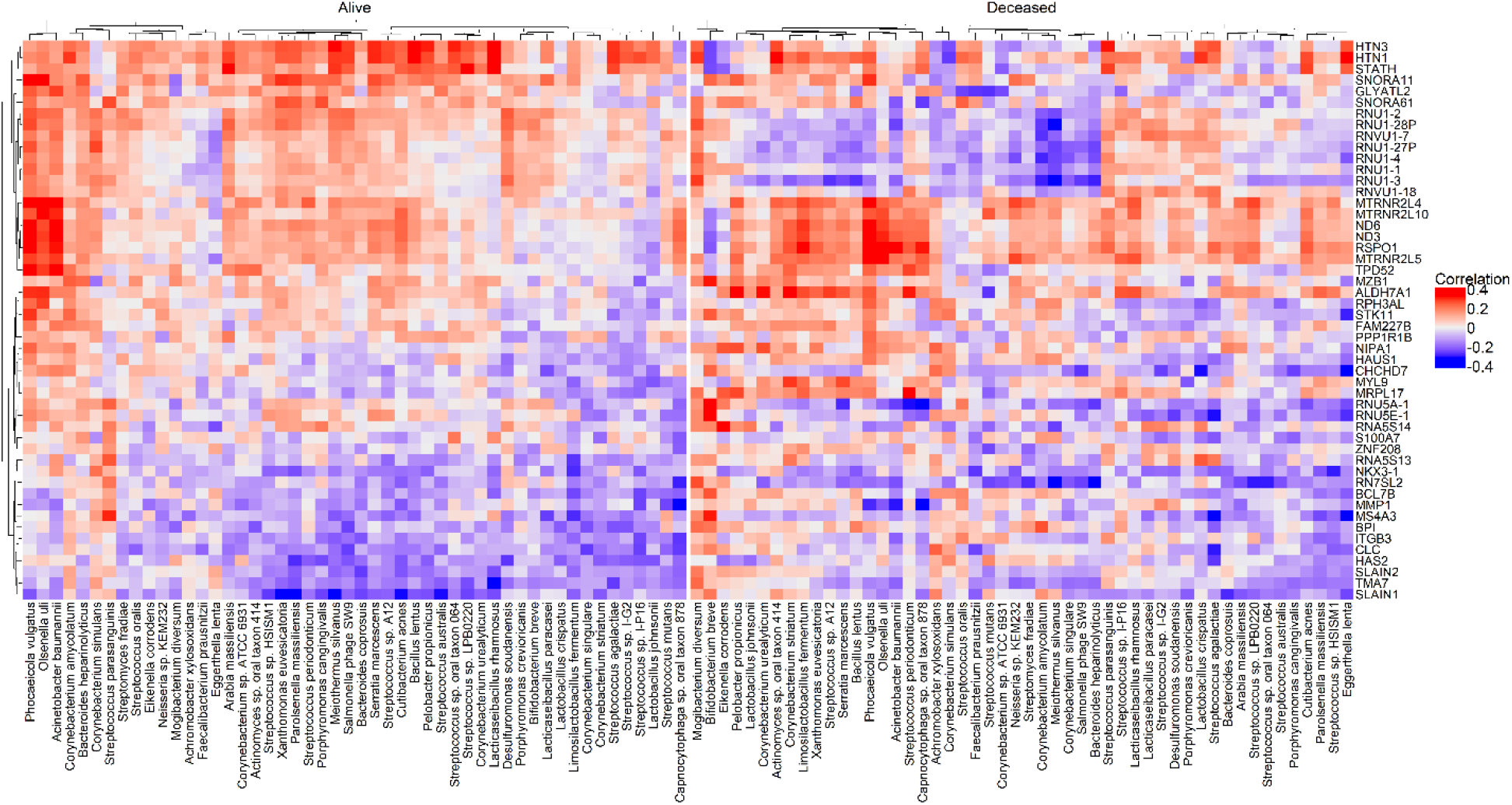
Heatmaps of Spearman’s rank correlations between the top 50 taxa from metagenome and the top 50 genes from host transcriptome, in alive and deceased groups, separately. The top 50 features were selected based on the proportion of selection in all CV iterations.

Table S6 reports the included features in the metagenomic, metatranscriptomic, and host transcriptomic risk scores separately. The feature importance was determined by the selection proportion among all CV iterations. For the host transcriptomic data, the fold change between deceased and alive was used to determine the feature importance when the selection proportions were the same. Here we take the top 50 features in each data as an illustration to investigate the correlation networks among these three datasets. Figures 7, S3, and S4 show the paired correlation heatmaps among the selected metagenomic, metatranscriptomic, and host transcriptomic features in the alive and deceased groups, respectively. Notably, the alive and deceased groups have different correlation patterns among these top 50 features from any two datasets. Specifically, the metagenomics features tend to have stronger correlations with the host transcriptomic and metatranscriptomic features in the deceased group, compared to the alive group; and the metatranscriptomic features tend to have more negative correlations with the host transcriptome in the alive group.

Note that the results reported in this section are different from those in Sulaiman et al. [26] in which the main goal was to reveal the scientific findings and the Cox proportional-hazards model [51] was employed to identify the candidate taxa and genes associated with the time to death. In this paper, we formally introduce the MRS concept and propose it as a general method with the detailed instruction on how to construct MRS. As a validation of the proposed method, the results presented above based on the binary outcome (Deceased vs. Alive) agree with the previous scientific conclusions [26]. Table 2 reports the hazard ratios of all risk scores constructed in this paper and their combinations on the time to death based on the Cox proportional-hazards model. All risk scores are significantly associated with the time to death. As we found in [26], metatranscriptomic data alone, or combined with the other two datasets, always has a higher hazard of death, because it involves SARS-CoV-2 viral, which is a key risk factor on the COVID-19 mortality.

**Table 2.**
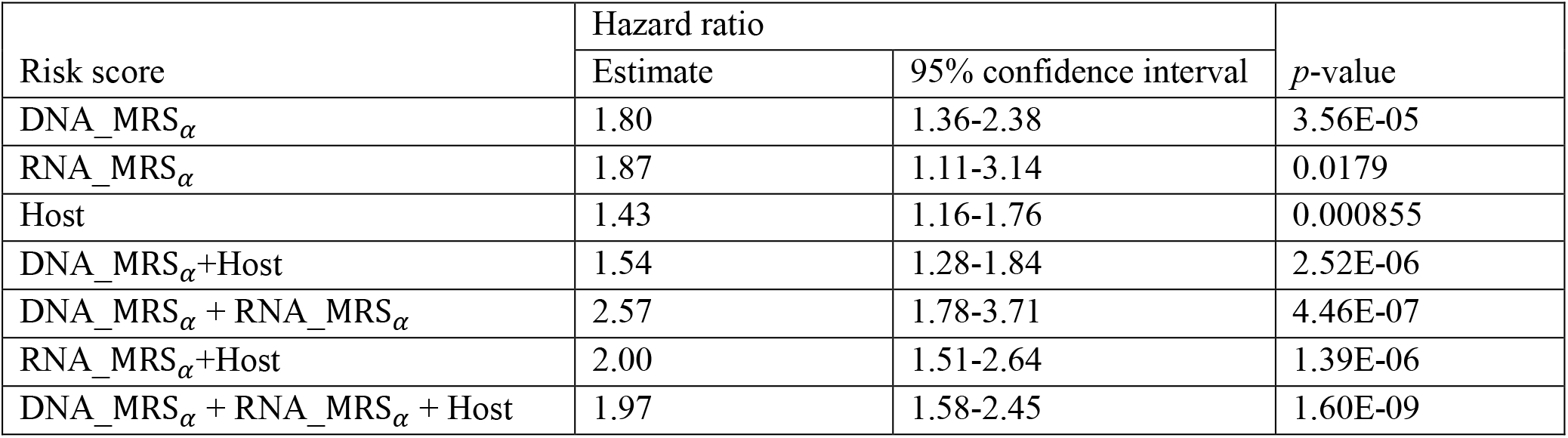
Association results between the risk scores and the time to death based on the Cox proportional-hazards model in the NYULH COVID-19 cohort.

| Risk score | Hazard ratio |  | *p*-value |
| --- | --- | --- | --- |
|  | Estimate | 95% confidence interval |  |
| DNA_MRS$_{\alpha}$ | 1.80 | 1.36-2.38 | 3.56E-05 |
| RNA_MRS$_{\alpha}$ | 1.87 | 1.11-3.14 | 0.0179 |
| Host | 1.43 | 1.16-1.76 | 0.000855 |
| DNA_MRS$_{\alpha}$+Host | 1.54 | 1.28-1.84 | 2.52E-06 |
| DNA_MRS$_{\alpha}$ + RNA_MRS$_{\alpha}$ | 2.57 | 1.78-3.71 | 4.46E-07 |
| RNA_MRS$_{\alpha}$+Host | 2.00 | 1.51-2.64 | 1.39E-06 |
| DNA_MRS$_{\alpha}$ + RNA_MRS$_{\alpha}$ + Host | 1.97 | 1.58-2.45 | 1.60E-09 |

Overall, these results highlight that the proposed community-based MRS_*α*_, can characterize and summarize the microbial profiles effectively and provide a flexible way to integrate microbiome data with other omics data. Integrations of risk scores from different omics data further improves the predictive performance on the alive/deceased status in the NYULH COVID-19 study.

#### TEDDY study

Although the TEDDY cohort includes both genome and microbiome data, the previous microbiome research on TEDDY study [53, 54] focused exclusively on the microbiome profiles and only identified very few microbial signatures associated with T1D. Given the fact that T1D is a multifactorial disease caused by both genetic and environmental factors and the children enrolled in the TEDDY study all have high genetic risk for T1D development (they have at least one of nine HLA DR-DQ genotypes associated with high risk for T1D) [29], we here propose a new angle to employ the proposed MRS along with the existing PRS for T1D to investigate the combined effect of microbial profile and host genetic profile on T1D risk prediction. Specifically, we analyzed 551 TEDDY subjects who have both microbiome data and genotype data; 75 of them developed T1D. Using the available genotype data and the PRS algorithm which has the robust and superior prediction performance on T1D [48, 49], we built the PRS for subjects. We used the microbial samples that were collected at the time point most close to month 30 when microbiome profile got stable and the largest sample size was available, to build MRS_*α*_ to predict T1D status independently. The practice of MRS calculations are the same as those used in the NYULH COVID-19 study.

Figure 8A compares the AUCs for predicting T1D based on the individual risk scores and the combination of PRS and MRS_*α*_, and Figures 8B-D show the Kaplan–Meier survival curve comparisons between high and low risk group identified by PRS, MRS_*α*_ and PRS + MRS_*α*_ respectively. Specifically, subjects whose risk scores are above the third quartile are defined as high risk, others as low risk. Although the predictive models considered in Figure 8A have only modest predictive ability in the TEDDY cohort (AUC range: [0.58, 0.63]), we found that integrating PRS and MRS_*α*_ scores is more useful in stratifying the subjects into high and low risk groups for T1D development (Figure 8D) than the PRS (Figure 8B) or MRS_*α*_ (Figure 8C) alone, which indicates that the potential genetic-microbial interaction effect on the T1D progression. These results exhibit the utility of modeling multi-omics risk scores to identify the high risk populations who can benefit from more targeted interventions.

**Figure 8.**
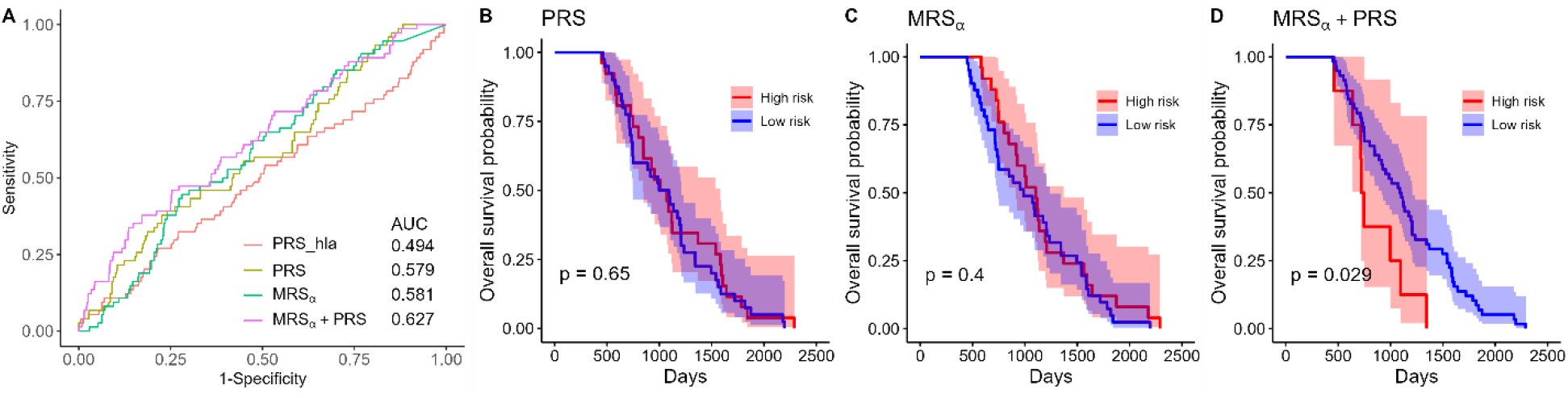
Results for T1D prediction in the TEDDY study. A. ROC curves and AUC values for predicting T1D status using various risk score. PRS_hla is constructed from the HLA alleles alone, and PRS is constructed from all SNPs found in the TEDDY cohort based on the existing PRS algorithm [49]. MRS_*α*_ is the negative alpha diversity (Shannon index) calculated on the selected sub-community, which is selected by ANCOM-BC method and P+T method. B–D. Kaplan–Meier plots for the groups of subjects at high and low risk of developing T1D, based on PRS, MRS_*α*_, and the combination of PRS and MRS_*α*_, respectively. Subjects whose risk scores are above the third quartile are defined as high risk, others as low risk, others as low risk.

## Discussion

With the recent proliferation of large-scale microbial association studies, we propose a two-step novel microbial risk score framework to aggregate the high-dimensional microbiome profile into a summarized risk score and apply it in disease prediction. Specifically, we first identify the associated taxa based on the recommended microbial association tests by two recent benchmarking works [32, 33] and P+T method, and then construct a community-based MRS_*α*_, because that the microbiome is a complex ecosystem composed of numerous sub-communities, and its influence on the disease development acts at the community instead of the single-microbe level and is disease-dependent. The application in the NYULH COVID-19 cohort demonstrates the superior performance of MRS_*α*_ in the disease prediction, compared to the standard MRS_*S*_, which is constructed similarly as PRS, ML-based prediction algorithms, and six alpha diversity measures on the whole microbiome community. The evaluation of MRS_*α*_ using the GMHI integrated dataset which consists of independent discovery and validation cohorts reveals the notable reproducibility of MRS_*α*_ in terms of disease prediction.

Combining omics datasets that provide biological information from different layers is vital to comprehensively study phenotypes and accurately predict diseases. However, complex data structures, for example, high-dimensionality, sparsity, compositionality, interdependence, and hierarchical tree structures, all make multi-omics data integration challenging. In this paper, the proposed MRS provides a straightforward and flexible way to incorporate multi-omics datasets and explore the microbial interactions with other omics profiles. Integration of the proposed MRS and the risk scores constructed from other omics data increases the ability for disease prediction. Integrations of metagenomic with metatranscriptomic and host transcriptomic datasets from NYULH COVID-19 cohort underline the critical and insightful utility of the constructed risk scores for disease prediction and the promising ability of multi-omics data integration for predictive accuracy improvement. Additionally, the data from TEDDY study illuminates the potential in combining MRS and PRS to explore genetic-microbial interaction and identify the high risk population.

Apart from the ANCOM-BC and Shannon index used in the proposed MRS_*α*_, there are other differential abundance methods available to identify the signature microbial taxa associated with disease and other alpha diversity indices to characterize the community diversity. Here we investigate how does using two other differential abundance methods (ALDEx2 [56] and Maaslin2 [57] suggested by [32, 33]) and Simpson and observed alpha diversity indices to construct MRS_*α*_ affect the predictive performance of the MRS framework in terms of AUC value and 95% CI in the discovery and validation cohorts [27] separately. Figure S5 shows that no single MRS_*α*_ can uniformly perform best for all predictions in the discovery and validation cohorts, as various DA methods have different model assumptions and test hypotheses and various alpha diversities indices have different definitions, while links between microbiome profile with various healthy or disease conditions are different. Specifically, given an alpha diversity index in the second step, DA method has no effect on the prediction performance of MRS_*α*_ in both discovery and validation cohorts (*p*-value>0.05) using the Kruskal-Wallis test on the AUC, except for Simpson index in the discovery cohort (*p*-value=0.03) (Figure S6). MRS_*α*_ s constructed with ANCOM-BC, ALDEx2, and Maaslin2, which all have been well-recognized [32, 33], have comparable performances. It supports our suggestion that to carry over the evaluation results of the DA tests from an objective benchmark work to guide the selection of DA test in the MRS framework. In terms of comparisons among Shannon, Simpson and Observed indices, Observed index based MRS_*α*_ has the highest AUCs, followed by Shannon index, while Simpson index has the lowest AUCs, in the discovery cohort (Figure S7). On the other hand, Shannon index consistently has better or comparable AUCs in the validation cohort. Meanwhile, Observed and Simpson indices introduce more variation in the predictive performance of MRS_*α*_ (Figure S7). Observed index lacks some reproducibility in the validation cohort, compared to its impressing performance in the discovery cohort, probably because it only accounts for species richness. Taken together, Shannon index based MRS_*α*_ has relatively more robust and consistent prediction performance. With existing discussions [32, 33] and the observations above in this manuscript, we include various DA methods commonly used and recommended in the microbiome association studies and various alpha diversity indices in the MRS R package to let the proposed MRS framework informative and more practically valuable.

The findings of this study have some limitations. First, considering microbial profile varies across ethnicities as well as geographies [58–60], it is necessary to evaluate the portability of MRS between populations. More advanced methods will be required to reduce the bias due to ethnical or geographical differences. Second, the microbiome data have versatile characteristics and unique features, such as phylogenetic tree structure, functional structure, hierarchical taxonomy, and dynamic nature, which also play critical roles in analytical accuracy and efficiency [61, 62]. Incorporating such features may improve the accuracy of MRS. Third, derivation and validation of MRS require large scale microbiome studies. However, the high cost of metagenomics sequencing restrict the comprehensive external validation.

Despite the above challenges, this paper proposes a practicable way to summarize the microbial profiles and provides promising findings for comprehensive microbiome research to bolster the microbiome’s utility as a potential source of novel therapeutic features.

## Conclusions

This paper sheds light on the utility of the microbiome data for disease prediction and multi-omics integration by converting the complex microbial profile into a continuous risk score. The proposed MRS tool provides great potential in studying the complex microbial ecosystem, understanding the microbiome’s role in disease diagnosis and prognosis, and exploring microbiome’s full clinical potential.

## List of abbreviations

ANCOM-BC: analysis of compositions of microbiomes with bias correction
AUC: area under the receiver operating characteristic curve
CA: colorectal adenoma
CC: colorectal cancer
CD: Crohn’s disease
CV: cross-validation
DA: differential abundance
DESeq2: differential expression analysis (v2)
GMHI: gut microbiome health index
GWASs: genome-wide association studies
HR: hazard ratio
MetaPhlAn: metagenomic phylogenetic analysis
ML: machine learning
MRS: microbial risk score
MWASs: microbiome-wide association studies
NYULH: NYU Langone Health
PRS: polygenic risk score
PD: phylogenetic diversity
P+T: Pruning and thresholding
QIIME: quantitative insights into microbial ecology
RA: rheumatoid arthritis
ROC: receiver operating characteristic
StrainPhlAn: metagenomic strain-level phylogenetic analysis
TEDDY: the environmental determinants of diabetes in the young
T1D: type 1 diabetes
T2D: type 2 diabetes (T2D)

## Declarations

### Ethics approval and consent to participate

All utilized microbiome datasets are publicly available. No ethics approval or consent to participate was required for this study.

### Consent for publication

Not applicable: All utilized microbiome datasets are publicly available. No consent for publication was required for this study.

### Availability of data and materials

For the NYULH COVID-19 cohort, all sequencing data used for this analysis are available in the NCBI Sequence Read Archive under project numbers PRJNA688510 and PRJNA687506 (RNA and DNA sequencing, respectively).

For the TEDDY study, TEDDY microbiome 16S rRNA gene sequencing data are publicly available in the NCBI database of Genotypes and Phenotypes (dbGaP) with the primary accession code phs001443. v1.p1, in accordance with the dbGaP controlled-access authorization process. Clinical metadata analysed during the current study will be made available in the NIDDK Central Repository at https://repository.niddk.nih.gov/studies/teddy/?query=teddy.

MRS R package used for the analyses is available at https://sites.google.com/site/huilinli09/software and https://github.com/chanw0/MRS, together with its manual. We also included the GMHI data and provided the code in the example section to reproduce the results in this manuscript.

### Authors’ contributions

CW developed the microbial risk score framework, performed data analyses, and wrote the manuscript. LS performed data analyses in the NYULH COVID-19 cohort and contributed to manuscript writing. JH performed data analyses in the TEDDY cohort and contributed to manuscript writing. BZ, RH, and JA contributed to the biological insights and interpretation, and to manuscript writing. HL contributed to the methodological ideas for the proposed framework, simulations, real data analyses, and manuscript writing. All authors read and approved the final manuscript.

## Acknowledgements

Not applicable.

## Additional material

### Additional file 1

**Figure S1.** The ROC curves and AUC values for various ML algorithms to predict the alive or deceased status in the NYULH COVID-19 cohort. A. Predication performance for elastic-net logistic regression (glmnet), penalized discriminant analysis (pda2), regularized random forest (RRF), and neural networks with feature extraction (pcaNNet) methods. B. Predication performance for naive Bayes (naïve_bayes), neural network (nnet), stochastic gradient boosting (gbm), and support vector machines with polynomial kernel (svmPoly) methods.

**Figure S2.** The AUC values and 95% CIs for MRS_*α*_s to classify healthy and nonhealthy and two disease conditions in the discovery and validation GMHI cohorts [27], respectively. CA: colorectal adenoma, CC: colorectal cancer, CD: Crohn’s disease, and RA: rheumatoid arthritis.

**Figure S3.** Heatmaps of Spearman’s rank correlations between the top 50 taxa from metagenome and the top 50 taxa from metatranscriptiome, in the alive and deceased groups, separately. The top 50 features were selected based on the proportion of selection in all CV iterations.

**Figure S4.** Heatmaps of Spearman’s rank correlations between the top 50 taxa from metatranscriptome and the top 50 genes from host transcriptome, in the alive and deceased groups, separately. The top 50 features were selected based on the proportion of selectin in all CV iterations.

**Figure S5.** Comparisons among various MRSs in terms of AUC value and 95% CI in the discovery and validation cohorts [27]. Here candidate taxa are identified by ANCOM-BC [31], ALDEx2 [56], and Maaslin2 [57], and the MRS_*α*_s are constructed by Shannon, Simpson, and Observed indices, respectively. DA: differential abundance, CA: colorectal adenoma, CC: colorectal cancer, CD: Crohn’s disease, and RA: rheumatoid arthritis.

**Figure S6.** The mean and standard derivation of the ranks of MRS_*α*_’s AUCs with ANCOM-BC, ALDEx2, and Maaslin2, respectively. For each alpha diversity index in each comparison of two diseases or healthy conditions, the AUCs of MRS_*α*_ with three DA methods were ranked 1-3. A higher rank represents a higher AUC. For each alpha diversity index, the Kruskal-Wallis test was performed to check difference among three DA methods. All: all samples were used for test. Statistical significance: ns: *p*-value>0.05; *: *p*-value≤ 0.05.

**Figure S7.** The mean and standard derivation of the ranks of MRS_*α*_’s AUCs with Shannon, Simpson, and Observed indices, respectively. For each DA method in each comparison of two diseases or healthy conditions, the AUCs of MRS_*α*_ with three indices were ranked 1-3. A higher rank represents a higher AUC. For each DA method, the Kruskal-Wallis test was performed to check difference among three alpha diversity indices. All: all samples were used for test. Statistical significance: ns: *p*-value >0.05; *: *p*-value ≤ 0.05; **: *p*-value ≤ 0.01; ***: *p*-value ≤ 0.001; ****: *p*-value ≤ 0.0001.

### Additional file 2

**Table S1.** Number of discovery and validation samples used for MRS evaluation and validation from the GMHI multi-study cohort.

**Table S2.** AUC values for six common alpha diversity indices on the whole community to predict alive and deceased status in the NYULH COVID-19 cohort.

**Table S3** The identified species for MRS_*α*_ construction in terms of comparisons among healthy, CA, CC, CD, RA, and nonhealthy based on the discovery samples in the GMHI multi-study cohort.

**Table S4.** Average and standard deviation of relative abundances of the identified species in Healthy, CA, CC, CD, and RA discovery samples from the GMHI multi-study cohort. The identified species are used for MRS_*α*_ construction in terms of pairwise comparisons of Healthy versus CA, CC, CD, and RA, respectively.

**Table S5.** Average and standard deviation of relative abundances of the identified species in Healthy, CA, CC, CD, and RA validation samples from the GMHI multi-study cohort. The identified species are used for MRS_*α*_ construction in terms of pairwise comparisons of Healthy versus CA, CC, CD, and RA, respectively.

**Table S6.** Factors used for metagenomic, metatranscriptomic and host transcriptomic risk scores in the NYULH COVID-19 cohort.

### Additional file 3

**Section S1** Computational details for risk scores

## References

1. Hu J, Koh H, He L, Liu M, Blaser MJ, Li H: A two-stage microbial association mapping framework with advanced FDR control. Microbiome 2018, 6(1):1–16.

2. Gilbert JA, Quinn RA, Debelius J, Xu ZZ, Morton J, Garg N, Jansson JK, Dorrestein PC, Knight R: Microbiome-wide association studies link dynamic microbial consortia to disease. Nature 2016, 535(7610):94–103.

3. Koh H, Livanos AE, Blaser MJ, Li H: A highly adaptive microbiome-based association test for survival traits. BMC genomics 2018, 19(1):1–13.

4. Koh H, Blaser MJ, Li H: A powerful microbiome-based association test and a microbial taxa discovery framework for comprehensive association mapping. Microbiome 2017, 5(1):1–15.

5. Ahn J, Sinha R, Pei Z, Dominianni C, Wu J, Shi J, Goedert JJ, Hayes RB, Yang L: Human gut microbiome and risk for colorectal cancer. Journal of the National Cancer Institute 2013, 105(24):1907–1911.

6. Kostic AD, Xavier RJ, Gevers D: The microbiome in inflammatory bowel disease: current status and the future ahead. Gastroenterology 2014, 146(6):1489–1499.

7. Hoffmann AR, Proctor L, Surette M, Suchodolski J: The microbiome: the trillions of microorganisms that maintain health and cause disease in humans and companion animals. Veterinary Pathology 2016, 53(1):10–21.

8. Kelly TN, Bazzano LA, Ajami NJ, He H, Zhao J, Petrosino JF, Correa A, He J: Gut microbiome associates with lifetime cardiovascular disease risk profile among bogalusa heart study participants. Circulation research 2016, 119(8):956–964.

9. Gilbert JA, Blaser MJ, Caporaso JG, Jansson JK, Lynch SV, Knight R: Current understanding of the human microbiome. Nature medicine 2018, 24(4):392–400.

10. Fattorusso A, Di Genova L, Dell’Isola GB, Mencaroni E, Esposito S: Autism spectrum disorders and the gut microbiota. Nutrients 2019, 11(3):521.

11. Integrative H, Proctor LM, Creasy HH, Fettweis JM, Lloyd-Price J, Mahurkar A, Zhou W, Buck GA, Snyder MP, Strauss III JF: The integrative human microbiome project. Nature 2019, 569(7758):641–648.

12. Wang C, Hu J, Blaser MJ, Li H: Estimating and testing the microbial causal mediation effect with high-dimensional and compositional microbiome data. Bioinformatics (Oxford, England) 2020, 36(2):347–355.

13. Bolyen E, Rideout JR, Dillon MR, Bokulich NA, Abnet CC, Al-Ghalith GA, Alexander H, Alm EJ, Arumugam M, Asnicar F et al: Reproducible, interactive, scalable and extensible microbiome data science using QIIME 2. Nat Biotechnol 2019, 37(8):852–857.

14. Truong DT, Franzosa EA, Tickle TL, Scholz M, Weingart G, Pasolli E, Tett A, Huttenhower C, Segata N: MetaPhlAn2 for enhanced metagenomic taxonomic profiling. Nature methods 2015, 12(10):902–903.

15. Truong DT, Tett A, Pasolli E, Huttenhower C, Segata N: Microbial strain-level population structure and genetic diversity from metagenomes. Genome research 2017, 27(4):626–638.

16. Choi SW, Mak TS-H, O’Reilly PF: Tutorial: a guide to performing polygenic risk score analyses. Nature Protocols 2020, 15(9):2759–2772.

17. Wand H, Lambert SA, Tamburro C, Iacocca MA, O’Sullivan JW, Sillari C, Kullo IJ, Rowley R, Dron JS, Brockman D: Improving reporting standards for polygenic scores in risk prediction studies. Nature 2021, 591(7849):211–219.

18. Turnbaugh PJ, Ley RE, Hamady M, Fraser-Liggett CM, Knight R, Gordon JI: The human microbiome project. Nature 2007, 449(7164):804–810.

19. McDonald D, Hyde E, Debelius JW, Morton JT, Gonzalez A, Ackermann G, Aksenov AA, Behsaz B, Brennan C, Chen Y: American gut: an open platform for citizen science microbiome research. Msystems 2018, 3(3):e00031–00018.

20. Xavier JB, Young VB, Skufca J, Ginty F, Testerman T, Pearson AT, Macklin P, Mitchell A, Shmulevich I, Xie L: The cancer microbiome: distinguishing direct and indirect effects requires a systemic view. Trends in cancer 2020, 6(3):192–204.

21. de Cárcer DA: A conceptual framework for the phylogenetically constrained assembly of microbial communities. Microbiome 2019, 7(1):1–11.

22. Coyte KZ, Rao C, Rakoff-Nahoum S, Foster KR: Ecological rules for the assembly of microbiome communities. PLoS biology 2021, 19(2):e3001116.

23. Cho I, Blaser MJ: The human microbiome: at the interface of health and disease. Nature Reviews Genetics 2012, 13(4):260–270.

24. Thukral AK: A review on measurement of Alpha diversity in biology. Agric Res *J* 2017, 54(1):1–10.

25. Whittaker RH: Evolution and measurement of species diversity. Taxon 1972, 21(2-3):213–251.

26. Sulaiman I, Chung M, Angel L, Tsay J-CJ, Wu BG, Yeung ST, Krolikowski K, Li Y, Duerr R, Schluger R et al: Microbial signatures in the lower airways of mechanically ventilated COVID-19 patients associated with poor clinical outcome. Nature Microbiology 2021, 6(10):1245–1258.

27. Gupta VK, Kim M, Bakshi U, Cunningham KY, Davis JM, Lazaridis KN, Nelson H, Chia N, Sung J: A predictive index for health status using species-level gut microbiome profiling. Nature communications 2020, 11(1):1–16.

28. Lee HS, Burkhardt BR, McLeod W, Smith S, Eberhard C, Lynch K, Hadley D, Rewers M, Simell O, She JX: Biomarker discovery study design for type 1 diabetes in The Environmental Determinants of Diabetes in the Young (TEDDY) study. Diabetes/metabolism research and reviews 2014, 30(5):424–434.

29. Rewers M, Hyöty H, Lernmark Å, Hagopian W, She J-X, Schatz D, Ziegler A-G, Toppari J, Akolkar B, Krischer J: The Environmental Determinants of Diabetes in the Young (TEDDY) study: 2018 update. Current diabetes reports 2018, 18(12):1–14.

30. Zheng P, Li Z, Zhou Z: Gut microbiome in type 1 diabetes: A comprehensive review. Diabetes/metabolism research and reviews 2018, 34(7):e3043.

31. Lin H, Peddada SD: Analysis of compositions of microbiomes with bias correction. Nature communications 2020, 11(1):1–11.

32. Nearing JT, Douglas GM, Hayes MG, MacDonald J, Desai DK, Allward N, Jones C, Wright RJ, Dhanani AS, Comeau AM: Microbiome differential abundance methods produce different results across 38 datasets. Nature Communications 2022, 13(1):1–16.

33. Lin H, Peddada SD: Analysis of microbial compositions: a review of normalization and differential abundance analysis. NPJ biofilms and microbiomes 2020, 6(1):1–13.

34. Wilcoxon F: Individual comparisons by ranking methods. In: Breakthroughs in statistics. Springer; 1992: 196–202.

35. Segata N, Izard J, Waldron L, Gevers D, Miropolsky L, Garrett WS, Huttenhower C: Metagenomic biomarker discovery and explanation. Genome biology 2011, 12(6):1–18.

36. Love MI, Huber W, Anders S: Moderated estimation of fold change and dispersion for RNA-seq data with DESeq2. Genome biology 2014, 15(12):1–21.

37. Robinson MD, McCarthy DJ, Smyth GK: edgeR: a Bioconductor package for differential expression analysis of digital gene expression data. Bioinformatics 2010, 26(1):139–140.

38. Mandal S, Van Treuren W, White RA, Eggesbø M, Knight R, Peddada SD: Analysis of composition of microbiomes: a novel method for studying microbial composition. Microbial ecology in health and disease 2015, 26(1):27663.

39. Kaul A, Mandal S, Davidov O, Peddada SD: Analysis of microbiome data in the presence of excess zeros. Frontiers in microbiology 2017, 8:2114.

40. Marcos-Zambrano LJ, Karaduzovic-Hadziabdic K, Loncar Turukalo T, Przymus P, Trajkovik V, Aasmets O, Berland M, Gruca A, Hasic J, Hron K: Applications of machine learning in human microbiome studies: a review on feature selection, biomarker identification, disease prediction and treatment. Frontiers in microbiology 2021, 12:313.

41. Gou W, Ling C-w, He Y, Jiang Z, Fu Y, Xu F, Miao Z, Sun T-y, Lin J-s, Zhu H-l: Interpretable machine learning framework reveals robust gut microbiome features associated with type 2 diabetes. Diabetes Care 2021, 44(2):358–366.

42. Ke G, Meng Q, Finley T, Wang T, Chen W, Ma W, Ye Q, Liu T-Y: Lightgbm: A highly efficient gradient boosting decision tree. Advances in neural information processing systems 2017, 30:3146–3154.

43. Vabalas A, Gowen E, Poliakoff E, Casson AJ: Machine learning algorithm validation with a limited sample size. PloS one 2019, 14(11):e0224365.

44. Lamri A, Mao S, Desai D, Gupta M, Paré G, Anand SS: Fine-tuning of Genome-Wide Polygenic Risk Scores and Prediction of Gestational Diabetes in South Asian Women. Scientific reports 2020, 10(1):1–9.

45. Jost L: Entropy and diversity. Oikos 2006, 113(2):363–375.

46. Gauthier J, Derome N: Evenness-Richness Scatter Plots: a Visual and Insightful Representation of Shannon Entropy Measurements for Ecological Community Analysis. Msphere 2021, 6(2):e01019–01020.

47. Blaser MJ: Missing microbes: how the overuse of antibiotics is fueling our modern plagues: Macmillan; 2014.

48. Padilla-Martínez F, Collin F, Kwasniewski M, Kretowski A: Systematic review of polygenic risk scores for type 1 and type 2 diabetes. International journal of molecular sciences 2020, 21(5):1703.

49. Perry DJ, Wasserfall CH, Oram RA, Williams MD, Posgai A, Muir AB, Haller MJ, Schatz DA, Wallet MA, Mathews CE: Application of a genetic risk score to racially diverse type 1 diabetes populations demonstrates the need for diversity in risk-modeling. Scientific reports 2018, 8(1):1–13.

50. Udler MS, McCarthy MI, Florez JC, Mahajan A: Genetic risk scores for diabetes diagnosis and precision medicine. Endocrine reviews 2019, 40(6):1500–1520.

51. Harrell FE: Cox proportional hazards regression model. In: Regression modeling strategies. Springer; 2015: 475–519.

52. Chatterjee N, Shi J, García-Closas M: Developing and evaluating polygenic risk prediction models for stratified disease prevention. Nat Rev Genet 2016, 17(7):392–406.

53. Vatanen T, Franzosa EA, Schwager R, Tripathi S, Arthur TD, Vehik K, Lernmark Å, Hagopian WA, Rewers MJ, She J-X: The human gut microbiome in early-onset type 1 diabetes from the TEDDY study. Nature 2018, 562(7728):589–594.

54. Stewart CJ, Ajami NJ, O’Brien JL, Hutchinson DS, Smith DP, Wong MC, Ross MC, Lloyd RE, Doddapaneni H, Metcalf GA: Temporal development of the gut microbiome in early childhood from the TEDDY study. Nature 2018, 562(7728):583–588.

55. Kuhn M: Building predictive models in R using the caret package. Journal of statistical software 2008, 28(1):1–26.

56. Gloor G: ALDEx2: ANOVA-Like Differential Expression tool for compositional data. ALDEX manual modular 2015, 20:1–11.

57. Mallick H, Rahnavard A, McIver LJ, Ma S, Zhang Y, Nguyen LH, Tickle TL, Weingart G, Ren B, Schwager EH: Multivariable association discovery in population-scale meta-omics studies. PLoS computational biology 2021, 17(11):e1009442.

58. Gaulke CA, Sharpton TJ: The influence of ethnicity and geography on human gut microbiome composition. Nature medicine 2018, 24(10):1495–1496.

59. Deschasaux M, Bouter KE, Prodan A, Levin E, Groen AK, Herrema H, Tremaroli V, Bakker GJ, Attaye I, Pinto-Sietsma S-J: Depicting the composition of gut microbiota in a population with varied ethnic origins but shared geography. Nature medicine 2018, 24(10):1526–1531.

60. He Y, Wu W, Zheng H-M, Li P, McDonald D, Sheng H-F, Chen M-X, Chen Z-H, Ji G-Y, Zheng Z-D-X: Regional variation limits applications of healthy gut microbiome reference ranges and disease models. Nature medicine 2018, 24(10):1532–1535.

61. Lozupone C, Knight R: UniFrac: a new phylogenetic method for comparing microbial communities. Appl Environ Microbiol 2005, 71(12):8228–8235.

62. Chen J, Bushman FD, Lewis JD, Wu GD, Li H: Structure-constrained sparse canonical correlation analysis with an application to microbiome data analysis. Biostatistics 2013, 14(2):244–258.

